# GIAnT: a Glutamate Imaging Analysis Toolbox

**DOI:** 10.64898/2026.08.07.743580

**Authors:** Michael E. Xie, Johannes Friedrich, Elizabeth Wirsching, Caleb Jones Shibu, Maedeh Seyedolmohadesin, Naveen Ouellette, Tim Wang, Karel Svoboda, Adam S. Charles, Kaspar Podgorski

## Abstract

Recent advances in fluorescent indicators and optical microscopy now enable *in vivo* synaptic imaging of glutamate, which transmits the majority of signals between neurons in the brain. Extracting fluorescence signals from these recordings is complicated by the minuscule scale and dense clustering of synapses on dendrites, as well as brain motion in behaving animals. Here we present the Glutamate Imaging Analysis Toolbox (GIAnT), a set of automated tools for glutamate imaging data that corrects sample motion, identifies active synapses with super-resolution precision, and extracts synaptic fluorescence signals. Compared to methods designed for cellular imaging, GIAnT reduces motion artifacts, more accurately identifies active synapses, and improves extracted signal quality by reducing contamination from overlapping synapses. By pairing *in vivo* glutamate imaging with *post hoc* expansion microscopy, we find that >70% of the putative synapses extracted using GIAnT matched one-to-one with glutamatergic synapses onto the postsynaptic cell. Our results establish GIAnT as an automated and validated pipeline for processing synaptic glutamate imaging data at scale.

## Introduction

Synapses, the structures where neurons connect and communicate with one another, underlie many critical aspects of neural function. Understanding the signals transmitted at synapses *in vivo* is vital to uncover how the brain processes information, how memories are formed, how connections change to support learning and adaptive behavior, and how these processes are affected by aging, injury, or disease.

Driven by recent development of genetically-encoded glutamate indicators, glutamate imaging has emerged as a promising method to directly record synaptic activity (Aggarwal et al. 2023; Aggarwal et al. 2026). Glutamate is released at ∼80% of synapses in the brain, triggered by action potentials in the presynaptic neuron, and is bound by closely apposed glutamate receptors on the post-synaptic neuron at millisecond timescales (Fonnum 1984). Glutamate imaging detects brief and localized increases in glutamate concentration to monitor synaptic transmission.

Measuring glutamate signals *in vivo* requires imaging at high spatial resolution and high speed, through living, light-scattering brain tissue. Two-photon microscopy can achieve these requirements (Denk & Svoboda 1997) and is commonly used for synaptic glutamate imaging (Y. Chen et al. 2026; Wright et al. 2025; W. Chen et al. 2024; Adoff et al. 2021). The resulting data are typically movies in which only one or a few neurons are labeled with a fluorescent glutamate indicator, and their dendrites or axons are recorded at high speed. In these movies, transient flashes indicate localized glutamate release. A vital step in processing these datasets is to identify the locations and release times of each imaged synapse. Automating this process is essential for robust, scalable, and reproducible analysis, but no validated pipeline for this currently exists.

For imaging of calcium indicators, there has been significant effort in establishing pipelines that automatically extract signals from fluorescence movies (Benisty et al. 2022; Charles et al. 2020). Current pipelines include motion correction (Pnevmatikakis & Giovannucci 2017; Hattori & Komiyama 2022), source extraction (Giovannucci et al. 2019; Pnevmatikakis et al. 2016; Stringer et al. 2026; Mishne et al. 2018; Charles et al. 2022), and validation (Gauthier et al. 2022; Song et al. 2021). These methods, however, are primarily designed and tested for signals at cell bodies (Giovannucci et al. 2019; Stringer et al. 2026), dendritic shaft and axons (Charles et al. 2022; Berlanga et al. 2025), or for widefield fluorescence (Charles et al. 2022; Saxena et al. 2020) and may not translate easily to tiny and densely clustered synapses.

Two-photon glutamate imaging presents challenges for image processing pipelines. First, imaging of glutamate indicators must be sufficiently fast (> 100 Hz) to capture the rapid fluorescence decay after a glutamate release event (on the order of 20 ms). To achieve a sufficiently high frame rate, two-photon glutamate imaging is typically performed in narrow strips that reduce the time per frame by collecting relatively few long lines scanned by a resonant scanner, which has a fixed line frequency regardless of scan amplitude. Analogous constraints apply to camera-based imaging, where readout speed is often determined by the number of image rows. Although strip imaging is highly practical, this approach leads to multiple challenges in conventional motion correction. For one, these images contain minimal anatomical features (the single dendritic branch) that can be used to register images across time. Moreover, typical sample motion on the order of several microns (Collman 2010) can easily move objects of interest out of the sampling window. Commonly-used tools developed for larger-scale neural imaging typically assume that the amplitude of motion is much smaller than the size of the imaging window (Pnevmatikakis & Giovannucci 2017; Hattori & Komiyama 2022). As this assumption does not hold for strip-imaging, these algorithms can fail when applied to such recordings. Second, source extraction is complicated by the small size of synapses and the limited resolution of optical imaging. Specifically, because synapses are small (<1 fL in volume) and dense along the dendrite (∼ 1–2 µm^−1^) (Ballesteros-Yáñez et al. 2006), glutamate imaging sources have spatial scales near the imaging diffraction limit and often overlap with each other in images. To distinguish these small neighboring sources, specialized localization methods are needed.

Prior methods to extract sources from dendritic glutamate imaging data have focused on manual labeling, finding source locations from correlation-based summary images, or extracting activity based primarily on anatomical structure (Wright et al. 2025; Adoff et al. 2021; Aggarwal et al. 2023; Yu et al. 2024). These methods, however, each face drawbacks. Manual annotation is time-intensive and subject to annotator bias. Graphical user interfaces (GUIs) and assistive programs (Berg et al. 2019; Z. Chen et al. 2025) can help with manual annotation of small datasets, but the sparsity of activity and long recording sequences make verifying both the location *and* timing of events untenable. Correlation-based summary images, such as nearest-neighbors correlation maps (Yu et al. 2024; Stringer et al. 2026), help reduce the data that needs to be analyzed to a single image, but are strongly limited by imaging resolution and may drown out sparsely active sources. Furthermore, none of these source extraction methods have been systematically validated for glutamate imaging data.

Here we present Glutamate Imaging Analysis Toolbox (GIAnT), an open source package for motion correction and source extraction for *in vivo* two-photon glutamate imaging. Our motion correction method, *StripRegistration*, uses a dynamically updated template to enable estimation of motion displacements larger than the imaged field of view. Our source extraction method, ***S****ource* ***I****dentification by Activity* ***Lo****calization* (SILo), uses superresolution spatial localization of glutamate transients to generate a summary image at higher resolution than correlation-based methods, resulting in better-resolved peaks for initializing segmentation algorithms. We validate the extracted sources through realistic simulations with statistics matched to real recordings and a novel paired *i*n vivo functional imaging and *ex vivo* histology dataset.

## Results

### GIAnT overview

GIAnT implements two main steps: motion correction and source extraction (Figure 1). Our MAT-LAB implementation of GIAnT is available on GitHub at https://github.com/AllenNeuralDynamics/GIAnT-MATLAB.

**Figure 1:**
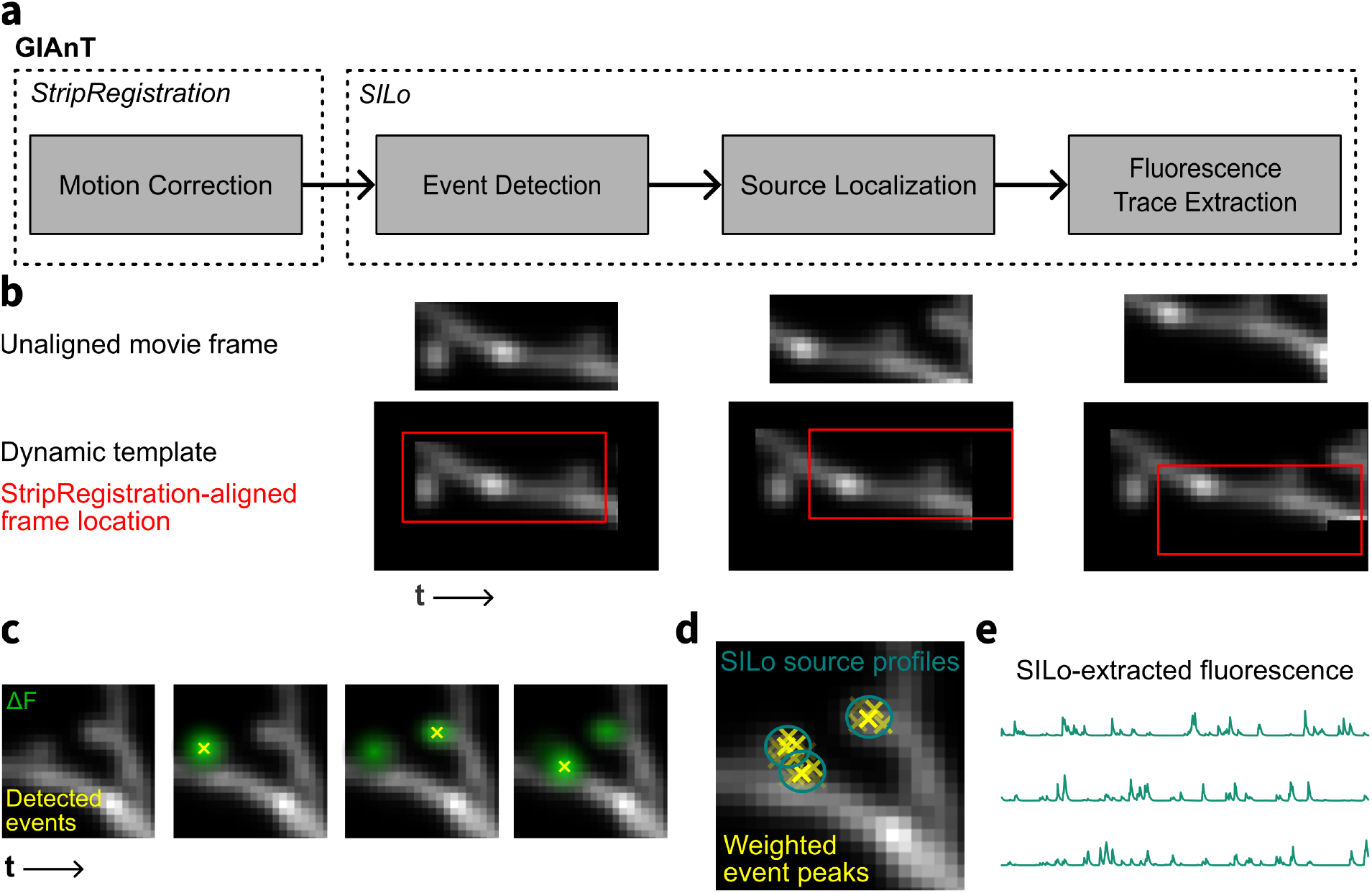
Schematic of Glutamate Imaging Analysis Toolbox (GIAnT). **a**. GIAnT consists of the StripRegistration motion correction algorithm, followed by Source Identification by Activity Localization (SILo). **b**. StripRegistration uses a dynamically updated template to improve alignment of large sample motion in strip-shaped fields of view. **c**. SILo starts with event detection, which is done by matched filtering the Δ*F* movie for the characteristic spatiotemporal shape of a glutamate release transient. **d**. To localize sources, detected event locations are weighted by their peak intensities and aggregated. Peaks of event density correspond to centers of SILo-detected source spatial profiles. **e**. A constrained matrix factorization is performed to extract fluorescence traces from the SILo-detected sources and refine their spatial profiles.

### Motion correction for recordings of narrow strips

To accurately and robustly correct for motion in strip-imaging data, we developed StripRegistration, an algorithm that allows for displacements larger than the size of the imaging window by iteratively building a template that extends beyond the bounds of the target imaging FOV (see **Methods**). An initial template is generated by aligning, using crosscorrelation, a subset of frames with high mutual correlation. The algorithm then aligns each raw data frame to the template in sequence. At each pixel in the image, the template is updated as an average of the aligned frames. Displacements can cause the template to become larger than the field of view size of the raw movie. To reduce outsized influence from undersampled pixels while retaining efficient discrete Fourier transform-based alignment, template pixels with fewer than a threshold number of observations are censored.

To evaluate StripRegistration, we ran the algorithm on both simulated and real data sets (Movie S1). We compared our approach to two popular rigid motion correction pipelines: NoRMCorre, as implemented in the calcium imaging processing software CaImAn (Pnevmatikakis & Giovannucci 2017; Giovannucci et al. 2019), and the motion registration algorithm implemented in Suite2p (Stringer et al. 2026). As the typical field of view for glutamate imaging of dendrites is small and the frame rate is high, non-rigid motion and rolling shutter effects are negligible, and thus we used only the rigid motion correction steps of each pipeline.

#### Simulation results

We first tested our motion correction algorithm on synthetic data simulated by taking an imaged volume of a dendrite expressing iGluSnFR4f and artificially 1) adding fluorescent flashes along the dendrite and spines 2) shifting the sampled field of view and plane at each frame, and 3) adding realistic noise (see **Methods**; Figure S1). We systematically varied the standard deviation of motion and overall signal-to-noise ratio by modulating the signal brightness, defined as the baseline photon count of the 99^th^ percentile-brightest pixel. These parameters enabled us to observe the motion correction behavior over a range of possible imaging conditions encountered *in vivo*.

On the simulated data, motion vectors estimated by StripRegistration had lower root mean squared error (RMSE) versus ground truth across the wide range of simulation motion amplitudes (Figure 2a), as well as across brightness levels up to 10× dimmer than typical *in vivo* recordings (Figure 2b). The error of predicted motion versus ground truth for frames that had large ground truth motion (>6 µm) was lower for StripRegistration compared to both Suite2p and CaImAn (Figure 2c).

**Figure 2:**
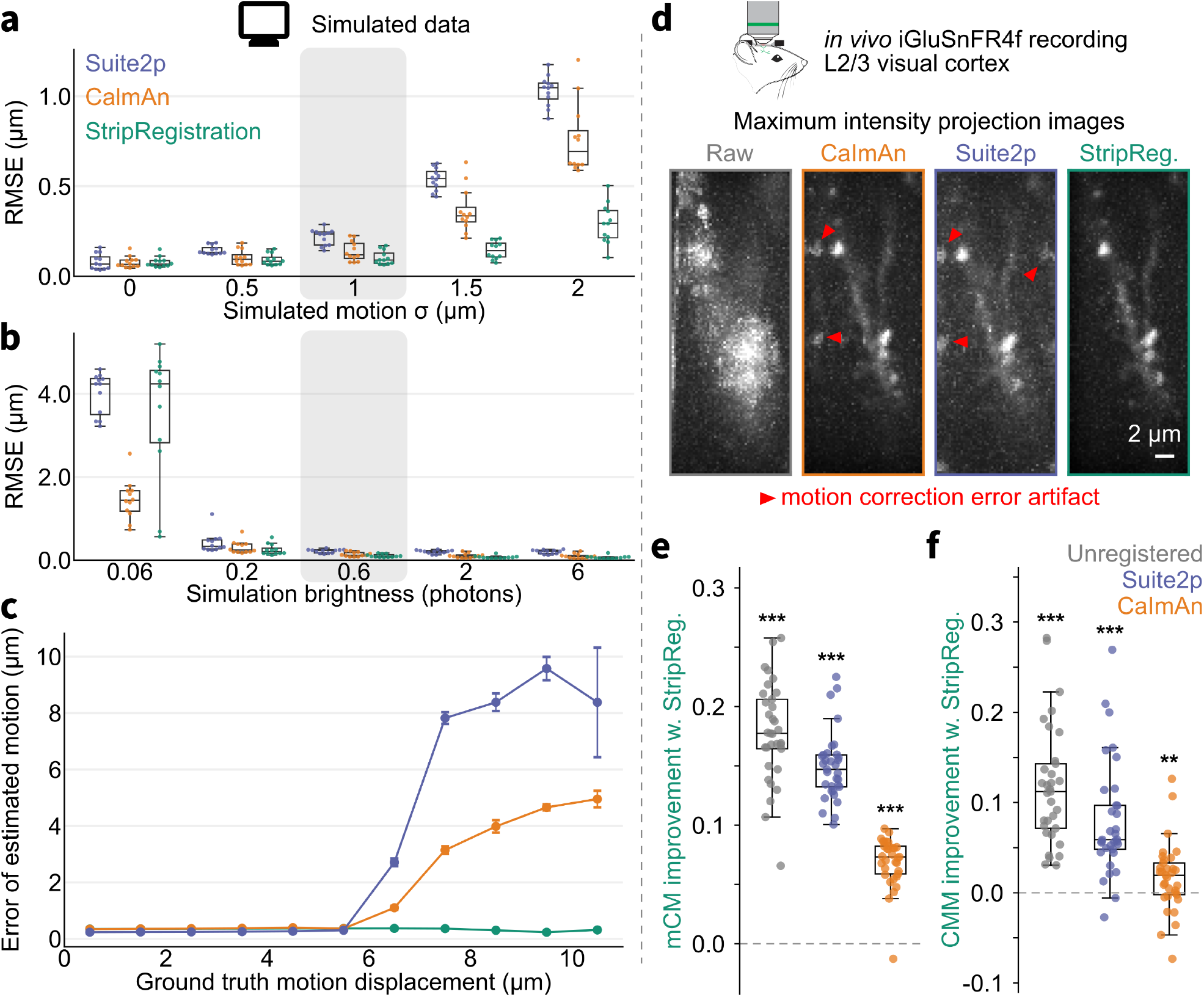
Evaluation of motion correction with StripRegistration. **a**. Root mean squared error (RMSE) between estimated and ground truth (GT) motion in simulated glutamate imaging of dendritic branches as a function of the standard deviation (σ) of simulated GT motion (simulation brightness fixed at 0.6 photons). **b**. RMSE between estimated and GT motion in simulated data as a function of simulated source brightness (motion σ fixed at 1 µm). In (a) and (b), grey boxes indicate the condition that most closely represents real data. **c**. Per-frame magnitude of the estimated motion vector error, as a function of the magnitude of simulated motion displacement in that frame (pooled over all frames in all simulations with brightness of 0.6 photons and motion σ of 1 µm). **d**. Maximum intensity projection of unregistered and registered *in vivo* data using the tested methods. Color scale is the same for all images. Data are *in vivo* two-photon recordings of iGluSnFR4f-expressing dendrites in L2/3 of visual cortex. Red arrows indicate artifacts in Suite2p and CaImAn due to incorrectly estimated large sample motion that are not present when using StripRegistration. **e**. Improvement of StripRegistration over other methods in mean frame correlation to the mean image (mCM). **f**. Improvement of StripRegistration over other methods in correlation of maximum projection image and mean projection image (CMM). In (e) and (f), *p<0.05, **p<0.01, ***p<0.001 on paired Wilcoxon signed-rank test with StripRegistration results.

#### Two-photon imaging results

We next assessed StripRegistration, alongside CaImAn and Suite2p motion correction, on *in vivo* two-photon recordings of dendrites expressing iGluSnFR4f in mouse visual cortex (see **Methods**). To evaluate residual motion artifacts, we first generated summary images consisting of the maximum intensity projections of each motion corrected video (Figure 2d). In the maximum intensity projection image, motion correction error artifacts get highlighted when dendritic structures (bright pixels) get incorrectly aligned to background regions (otherwise dim pixels). These artifacts and other motion correction errors can be quantified with two previously-established metrics: the mean correlation to the mean image (mCM) and correlation between the maximum projection image and mean image (CMM) (Hattori & Komiyama 2022). mCM measures how well *on average* individual frames are aligned to the mode of the data, and CMM measures how offset the worst-case frames are from each other. StripRegistration scored better on both metrics than either CaImAn and Suite2p (Figure 2e-f).

### Identifying sources in simulated glutamate imaging recordings

In glutamate imaging recordings, release events appear as localized flashes with a characteristic temporal exponential decay and small, circular spatial profile. In most brain areas, including cortex, presynaptic glutamate-releasing neurons exhibit low firing rates (0.1-10 Hz) (O’Connor et al. 2010; Niell & Stryker 2008) compared to the time constants of glutamate indicators (20-150 ms), resulting in flashes that appear sparsely in time and space. However, despite the sparsity of release events, synapses can be so close as to be unresolvable by the microscope (Ballesteros-Yáñez et al. 2006). To identify the locations of sources and disambiguate optically-overlapping synapses, we leverage the spatiotemporal sparsity of flashes to conduct super-resolution localization (Lelek et al. 2021), which precisely estimates the center of each flash (see **Methods**).

In the first step of SILo, we highpass filter each pixel and z-score the resulting intensities using time-varying estimates of baseline intensity and noise. We then apply a matched filter designed to detect release events, with a Difference-of-Gaussians profile in space—to capture localized fluorescence flashes—and exponential decay in time (see **Methods**). We detect putative events as local maxima in the filtered recording in space and time. We aggregate these maxima in quadrature over time to produce a single summary image we call the SILo activity image. The activity image provides a high-resolution representation of activity that localizes to the center of true sources more tightly than the conventional nearest-neighbors correlation (NNCorr) statistic (Yu et al. 2024; Giovannucci et al. 2019), as validated in simulations (Figure 3a-b). Peaks in the SILo activity image can then be used to seed a constrained non-negative matrix factorization that estimates the full spatial and temporal profiles of each source using their known centers.

**Figure 3:**
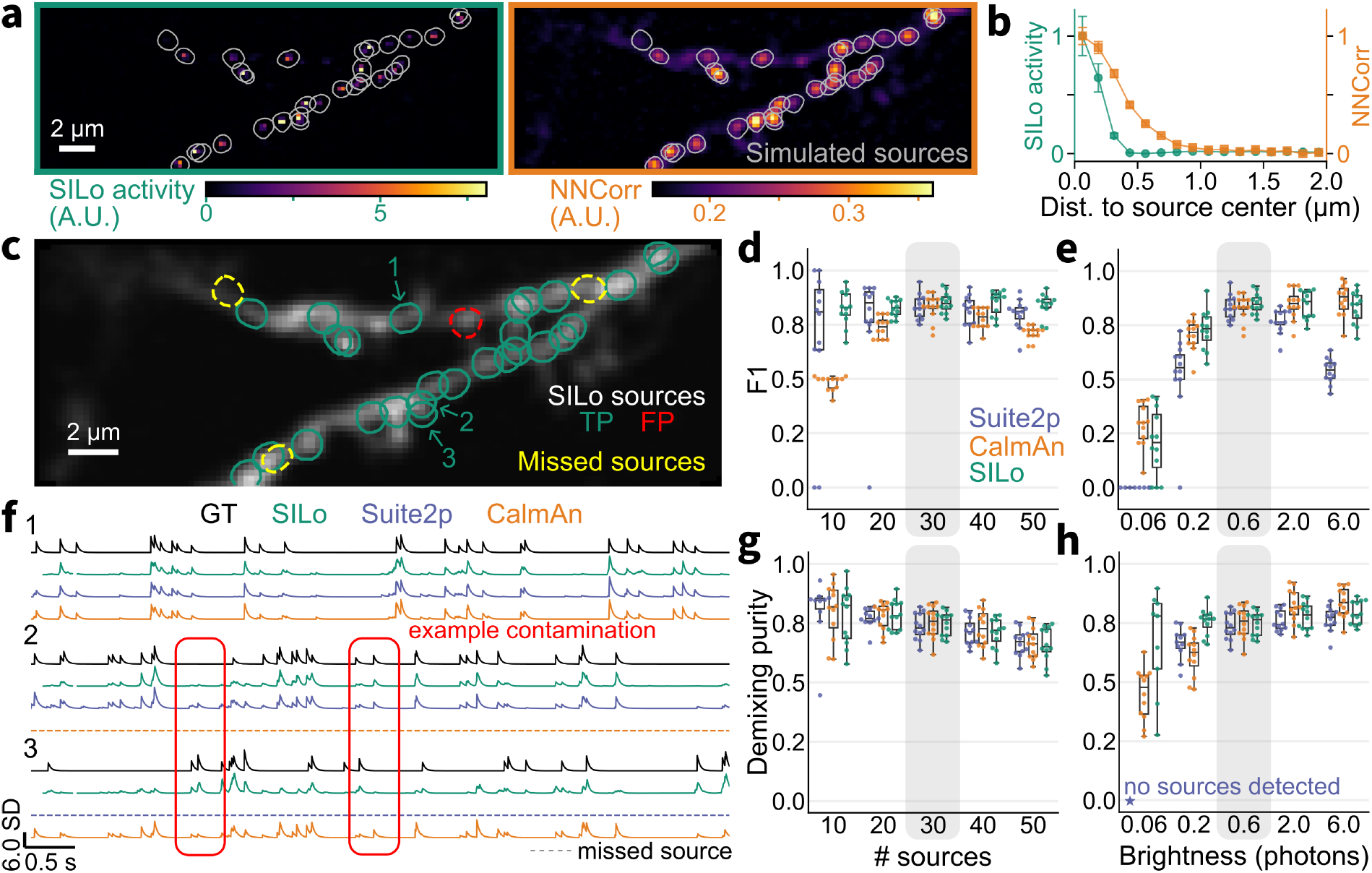
Validation of glutamate source extraction with simulations. **a**. Super-resolution activity image computed by SILo (left) and nearest neighbors correlation (NNCorr) image (right) with overlaid spatial profile contours of simulated sources (grey). **b**. Average activity metric value (normalized) by distance of pixel from the nearest ground truth source centroid. **c**. Contours of sources identified by SILo colored by true positive (TP) and false positive (FP). Contours of GT sources that were not detected by SILo are in yellow. **d**. F1 score of identified sources for each method compared to GT across simulations where the number of simulated sources is varied (brightness fixed at 0.6 photons). **e**. F1 score of identified sources for each method compared to GT across simulations where brightness is varied (number of sources fixed at 30). **f**. Example extracted traces of the three sources labeled in (c). Dotted lines indicate that the source was not detected by that method. Red boxes shows a time window where simulated activity from source 3 can contaminate the extracted activity from source 2. **g**. Average demixing purity of extracted TP traces for each method across simulations where the number of simulated sources is varied (brightness fixed at 0.6 photons). **h**. Average demixing purity of extracted TP traces for each method across simulations where brightness is varied (number of sources fixed at 30). Only TP sources with > 1 overlapping GT source were included for (g) and (h). In (d), (e), (g), and (h), grey boxes indicate the “default” simulation condition which most closely represents real data and was used to optimize parameters for each method.

We first validated that SILo was able to faithfully recover ground truth sources using the same simulations used to evaluate motion correction (Figure 3c) and compared its performance with other automated source extraction methods (CaImAn and Suite2p). In these simulations, we varied signal brightness and the number of generated active sources. To ensure fair comparison, we used the StripRegistration motion corrected movie as input to all algorithms, made modifications to the existing algorithms to avoid errors due to data lost during large movements, and systematically optimized the tunable parameters for all methods to maximize F1 score, using a training dataset with statistics matched to real recordings (see **Methods**). During this optimization, we found that Suite2p’s *sparsery* algorithm consistently performed better than its *sourcery* algorithm, so we used Suite2p with *sparsery* throughout. Source identification on simulations with the same parameters used for optimization were similar across methods, but SILo was more stable across simulation conditions than Suite2p and CaImAn (Figure 3d-e). Aggregating across all conditions, mean±s.e.m. F1 scores for SILo were 0.759±0.021, higher than for CaImAn and Suite2p (Caiman: 0.694±0.018, *p* < 0.001; Suite2p: 0.637±0.028, *p* < 0.001; paired Wilcoxon signed-rank tests).

We next evaluated the activity traces extracted by the three methods (Figure 3f). For each extracted source, we calculated *demixing purity* (see **Methods**), which reflects the ground truth source’s contribution to the extracted trace as a fraction of the total contribution from all spatially overlapping ground truth sources. The average demixing purity across sources in a dataset was more stable over a range of simulated source densities and brightnesses for SILo compared to both Suite2p and CaImAn (Figure 3g-h). Aggregating across all conditions, the average demixing purity for SILo was 0.745±0.010 (mean±s.e.m.), CaImAn was 0.717±0.014, and Suite2p was 0.735±0.008. SILo had a significantly higher demixing purity compared to Suite2p (*p* < 0.001, paired Wilcoxon signed-rank test) but not compared to CaImAn (*p* = 0.734, paired Wilcoxon signed-rank test).

### Extracting sources from in vivo glutamate imaging recordings

We next evalutated our processing methods using real data from *in vivo* glutamate imaging experiments. We imaged iGluSnFR4f-expressing dendrites in layer 2/3 of visual cortex of awake mice while presenting drifting grating stimuli (Figure 4a). In each trial, eight stimulus directions were randomly shuffled and presented for 2 s each, with 1 s of uniform luminance presented in between each grating presentation. We processed recordings using StripRegistration and SILo.

**Figure 4:**
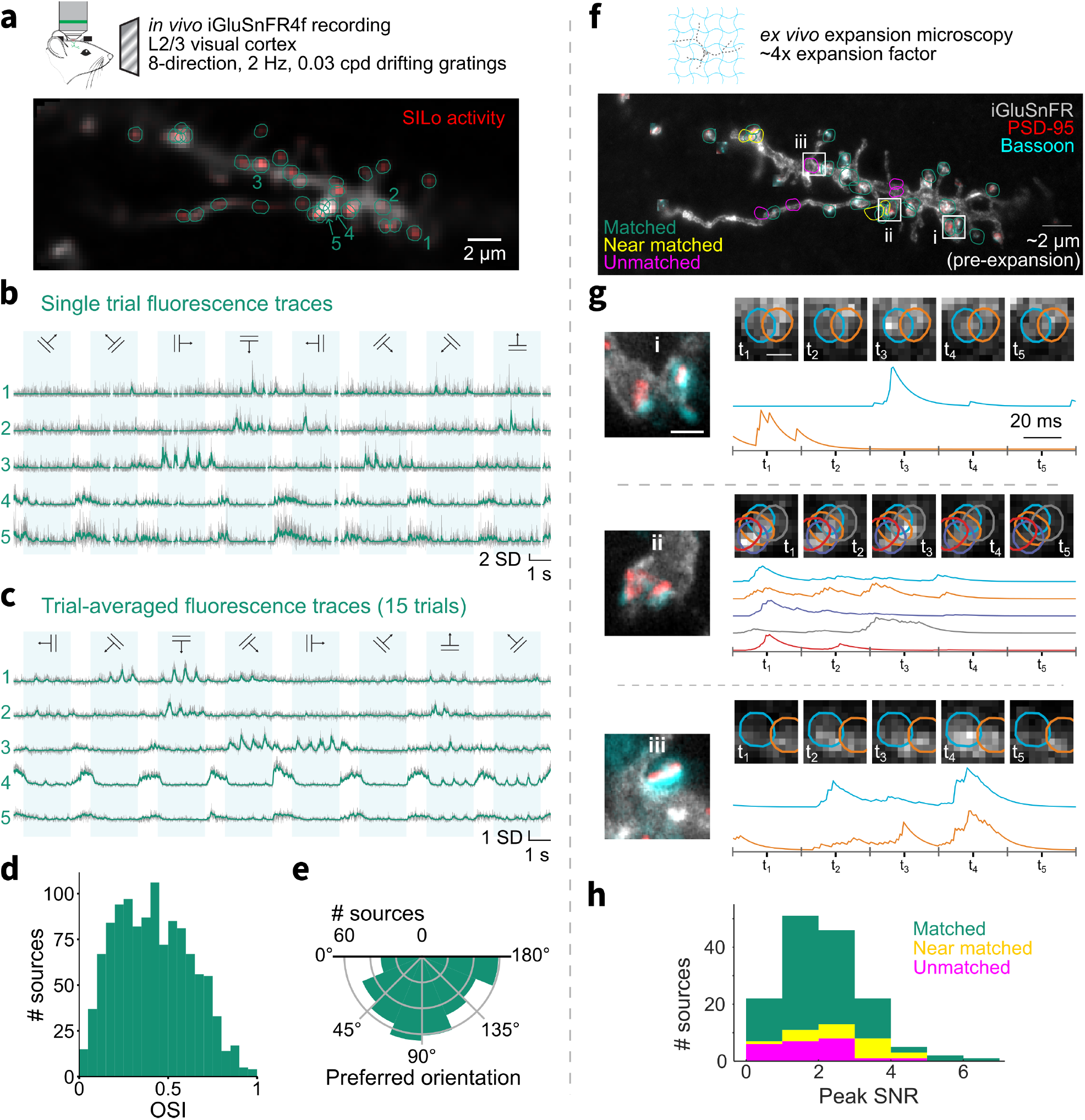
Characterization of sources extracted by SILo from *in vivo* data. **a**. iGluSnFR4f-expressing dendrites were imaged *in vivo* using two-photon microscopy while the mouse watched drifting gratings (8 directions, 2 Hz temporal frequency, 0.03 cycles/degree). Mean image of the imaged dendrite with overlaid SILo activity image and 1/*e*-level contours of identified sources are shown. **b**. Activity of selected sources (denoised estimate in green, least squares estimate in gray) during a single trial. Blue boxes denote when a drifting grating stimulus was presented. **c**. Trial-averaged activity of selected sources by direction of drifting grating. Blue boxes again denote when the drifting gratings were presented. **d**. Distribution of orientation selectivity indices (OSIs) of 1158 sources extracted from 31 recordings. **e**. Distribution of preferred orientations of the 417 orientation selective sources (OSI≥0.5) **f**. Maximum projection image of immunolabeled iGluSnFR and annotated areas of immunolabeled PSD-95 and Bassoon. PSD-95 and Bassoon channels are cropped to a small window around each manual synapse annotation to reduce visual clutter of synapses that are not on the iGluSnFR-expressing dendrite. The full histology volume across all channels is in Movie S2. 1/*e*-level contours of SILo-detected sources are aligned and overlaid, classified as matched, nearly matched (within 0.75 µm), and unmatched to a histology synapse **g (left)**. Cropped areas of iGluSnFR-labeled dendrite and PSD-95 and Bassoon labels in a plane with labeled dendrite or spines that are within the white boxes in (f). Scale bar, 0.5 µm. **g (right)**. Example frames from *in vivo* two-photon imaging corresponding to the synapses and sources around the areas on the left. Overlaid 1/*e*^2^-level contours show the distinct SILo-identified sources. Below the frames are corresponding extracted fluorescence, with fluorescence displayed in arbitrary units. The frames above the traces are generated by averaging over the corresponding time windows indicated in gray on the right. Scale bar (space), 1 µm. **h**. Stacked bar plot of peak signal-to-noise ratio (SNR) for matched, near matched, and unmatched sources.

Extracted sources (Figure 4a-b) showed tuning to specific orientations or directions (Figure 4c), which is characteristic of inputs to pyramidal neurons in visual cortex (Niell & Stryker 2008). The median orientation selectivity index (OSI) of the extracted sources across all datasets was 0.412 (IQR: 0.246–0.579, Figure 4d). Orientation selective sources (OSI> 0.5) showed preferences spanning all possible orientations (Figure 4e).

### Matching functionally-identified sources to histologically-identified synapses

We next sought to determine whether the glutamate sources identified using GIAnT correlated to synapses detected ultrastructurally. The same day after the iGluSnFR-labeled dendrites were imaged *in vivo*, the mouse was perfused (see **Methods**). Its brain was then sliced parallel to the cranial window and processed for expansion microscopy (F. Chen et al. 2015) and immunofluorescence to resolve sub-synaptic structures *ex vivo*. We immunolabeled Bassoon, a scaffolding protein located at pre-synaptic active zones, and PSD-95, a scaffolding protein located at the post-synaptic density of excitatory synapses; to identify the branch dendrite that was imaged *in vivo*, we also immunolabeled iGluSnFR (Figure 4e, Movie S2). We obtained paired function and histology datasets for four mice.

We aligned the time-averaged *in vivo* structure with the iGluSnFR channel in the histology image and manually annotated excitatory synapses in the *ex vivo* images for five dendrite branches. Synapses were defined as sites of iGluSnFR-labeled membrane with apposed bright spots in both the PSD-95 and Bassoon channels. When the sources extracted from SILo were matched one-to-one with histologically-identified synapses (Figure 4f), we found that SILo had a positive predictive value (PPV) for synapses in histology of 0.73 ± 0.04 (mean ± s.e.m. across five data sets, see **Methods**). Among the SILo sources that did not match uniquely with a histologically-identified synapse, we found that 45% were near matches, defined as being less than 0.75 µm away from an annotated synapse that was matched to another source. This indicates splitting of individual histologically-defined synapses into multiple sources by SILo, which could reflect errors in source extraction, uncertainty in histological annotation, or to the presence of multiple distinct release sites within a single large synapse. Of the 149 SILo sources identified across the five annotated histology datasets, 23 (15.4%) were not near a histologically-identified synapses on the imaged neuron. Sources that had extracted fluorescence traces with lower peak signal-to-noise ratio (SNR) were more likely to be unmatched with a histology synapse compared to sources with higher peak SNR (Figure 4i). Of the extracted sources with peak SNR less than 3, 21 (18.3%) were unmatched, whereas of the sources with peak SNR greater than 3, only 2 (5.9%) were unmatched.

## Discussion

Glutamate imaging has been rapidly adopted as a versatile method for studies of synaptic physiology, neuronal computation, circuit dynamics, and disease mechanisms. These emerging applications necessitate a complementary effort to validate and characterize different methods for extracting synaptic signals from these datasets.

In this study, we introduced a new analysis pipeline for glutamate imaging recordings, GIAnT, as well as methods and data for systematically evaluating pipelines against each other. The validation data we generated included both realistic simulations of glutamate imaging and correlative *in vivo* two-photon glutamate recordings with post-hoc *ex vivo* expansion microscopy of the same dendrites.

We compared GIAnT to the Suite2p and CaImAn data analysis pipelines, after optimizing the tunable parameters for all methods to suit glutamate recordings. StripRegistration outperformed other methods on the strip-shaped FOVs best suited for fast glutamate indicators, under motion conditions encountered *in vivo*. Robust motion correction is critical as errors in motion correction will either cause artifacts in downstream source extraction or necessitate strict censoring of frames, leading to inefficient use of data. Our method expands a dynamic template in space as frames are processed sequentially, which achieves excellent performance and is in principle compatible with online approaches. Alternatively, decentralized motion correction methods demonstrated for high-density extracellular recordings (Windolf et al. 2025; Varol et al. 2021) could also be adapted for these data to provide more contextual information early in the recording.

SILo matched or exceeded the performance of other methods for source extraction, and was especially robust to variations in the brightness and density of simulated sources. These variables can be unknown in real data and vary across recordings for multiple reasons, making this robustness especially valuable for performing *in vivo* experiments at scale. More accurate source detection also led to improved demixing of overlapping synapses in the fluorescence traces, especially in challenging simulations with high noise and dense synapses.

There also exist pipelines specifically for extracting sources from two-photon functional synaptic imaging data (Yu et al. 2024), but we do not focus on these as benchmarks because they identify source locations primarily based on anatomical structure. As some sources of glutamate indicator fluorescence are not possible to resolve using structural data alone (i.e. spines above and below the shaft, closely neighboring spines, or shaft synapses), use of the full spatiotemporal information is critical to reduce undetected true sources. However, information about the anatomy of the dendrite could in principle be integrated with our source extraction method to provide complementary information about the geometry of structures associated with the source (Z. Chen et al. 2025; Bernal-Garcia et al. 2025).

Superresolution localization of activity has previously been applied to identify sources in one-photon voltage imaging recordings (T.-W. Chen et al. 2025). Both approaches exploit the temporal sparsity of these transients to precisely identify sources whose spatial profiles overlap, but there are important differences that necessitate modifications to optimize performance for glutamate imaging data. Specifically, in SILo, we filter for the characteristic temporal transient shape of the imaged glutamate indicator and use an estimate of the point spread function as a spatial template, as synapses can be modeled as diffraction-limited points.

Lastly, for validation, we developed a set of realistic simulations of *in vivo* glutamate imaging data and collected a set of *in vivo* glutamate imaging data with retrospective histological ground truth. These validation datasets will also enable future iteration and optimization of data analysis pipelines for glutamate imaging. Validation data for calcium imaging has been widely adopted and have led to more systematic methods of improving corresponding analysis algorithms (Song et al. 2021), and our present study will help enable that same process for the field of glutamate imaging. In our histology dataset, we found some complex synaptic structures that had large or multiple areas of pre- and post-synaptic marker interaction. For such regions, it was difficult to ascertain how many distinct sources of glutamate release were present. In the future, further developments in expansion microscopy—for example to increase the expansion factor or label all proteins (Tavakoli et al. 2025; Damstra et al. 2022; M’Saad & Bewersdorf 2020)—could clarify whether these complex synaptic structures correspond to a single presynaptic excitatory neuron or multiple.

An additional benefit of GIAnT is that it can generalize well to imaging modalities that do not construct conventional images, such as band scanning in random access projection microscopy (Podgorski et al. 2023). Using SILo for source detection only requires an approximate point spread function of the microscope, which can be empirically measured for any imaging system. Filtering for sources can be performed even with sparse measurements, by appropriately normalizing the spatiotemporal convolutions. These new imaging modalities are particularly promising methods for widely sampling a neuron’s dendritic arbor in a single, simultaneous recording and at sufficiently high rates.

As imaging technologies advance in speed and scope—offering the ability to study *in vivo* synaptic activity at scale—automated pipelines, such as GIAnT, will become vital for effective analysis of these datasets.

## Methods

### Animal care and use statement

Experimental procedures involving mice were performed in accordance with National Institutes of Health (NIH) guidelines and approved by the Allen Institute Institutional Animal Care and Use Committee (IACUC) under protocols # 2415 and # 2109.

### In vivo two-photon glutamate imaging in mouse visual cortex

Mice of around nine weeks of age were implanted with a headplate for imaging of iGluSnFR4f. To achieve sparse labeling of neurons, each mouse was injected with a viral mix of 2E12 AAV1-hSyn-FLEX-iGluSnFR4f and 1.2E8 AAV9-CamKII-Cre in the visual cortex (VISp; 300 µm deep).

Mice were imaged at least 6 weeks after undergoing viral injection and headplate implantation. Neurons were imaged using a commercial 12 KHz resonant scanning microscope (Bergamo II, Thorlabs) controlled by ScanImage (MBF Bioscience). A Olympus 25x 1.0 NA objective equipped with a spherical aberration correction collar was used. A 1030nm laser (InSight X3, Newport) was used for two-photon excitation and emitted fluorescence was detected with a GaAsP photomutiplier tubethrough a 525/50 band-pass filter (Semrock). Dendrites of neurons expressing iGluSnFR4f were imaged at around 440 Hz, using 50 lines by 128 pixels per line at 10x software zoom. Mice were headfixed and awake during the experiment and passively viewing drifting gratings of varying orientations in 45^◦^ increments and with a spatial frequency of 0.03 cycles/degree and temporal frequency of 2 Hz. Each grating was presented for 2 s and uniform luminance was presented for 1 s between gratings.

Volumetric stacks of 21 planes spaced 0.5 µm apart and centered on the dendrite of interest were also collected.

### Simulated glutamate imaging datasets

Simulated glutamate imaging movies, with a field of view size of 125×45 pixels and pixel size of 250 nm, were created from volumetric stacks collected during the *in vivo* two-photon imaging experiments. The volume was scaled such that the 99th percentile pixel value equaled a specified brightness level. This brightness level was systematically varied across simulations (0.06, 0.2, 0.6, 2.0, and 6.0 photons).

The locations of a specified number of sources were selected by randomly sampling from bright voxels in the stack. The number of sources was systematically varied across simulations (10, 20, 30, 40, and 50 sources). Small subvoxel jitter was randomly added to the source locations and sources were selected such that they are at least 0.5 voxels from its nearest source. The shape of each source was simulated as a Gaussian of standard deviation 0.33 µm multiplied by the underlying anatomical structure around the source. The activity corresponding to each source was generated by simulating independent time-varying spike rates for each source. The presence of a spike for a source on each frame was sampled from a Bernoulli with a spike probability corresponding to the spike rate on that frame. As the spike probabilities on each frame was small, this approximately simulated a Poisson process. The Δ*F*/*F* amplitude of each generated spike was sampled from a log normal distribution (µ = 0 and σ = 0.25 for the underlying normal distribution). To simulate glutamate indicator fluorescence transients, the spike train was then convolved with a decaying exponential kernel with decay rate of 27 ms.

Sample motion was simulated by a smoothly time-varying three-dimensional vector, with correlations between the X, Y, and Z displacements modeled on *in vivo* observations. The motion displacement was centered at the origin and scaled to a specified standard deviation. This motion standard deviation was systematically varied across simulations (0, 0.5, 1, 1.5, and 2 µm).

Frames were simulated by sampling the volume, shifted by the corresponding simulated motion displacement. The sample was multiplied by an exponentially-decaying (τ = 1200 s) photobleaching curve. A frametime of 2.3 ms (∼ 435 Hz imaging rate) was used to sample in time. Poisson shot noise, log-normal detector noise, and Gaussian electronic noise were simulated independently for each pixel and frame on the observed planes.

Simulations with a motion standard deviation of 1 µm, brightness of 0.6 photons and 30 sources were denoted as “default” simulations as they were most similar to real recordings (Figure S1). These simulations were used for optimization of analysis pipeline parameters.

### Analysis of glutamate imaging datasets

#### Motion correction with StripRegistration

StripRegistration is a cross-correlation-based method. To help reduce the effect of noisy measurements, the raw movie can be downsampled by averaging every 2^*d*^ frames, where *d* is the specified downsampling factor parameter `ds_time`. If downsampling is used, all the following time points or frames mentioned are in the downsampled temporal space, unless otherwise specified.

First, an initial StripRegistration template is generated by registering the first 1000 frames using NoRMCorre (Pnevmatikakis & Giovannucci 2017). In order to maximize the performance of NoRMCorre, it is provided with a template that is the mean of a cluster of highly correlated frames within the initial 1000 frames. The average of the NoRMCorre-aligned 1000 frames is then padded evenly on all sides with NaNs so the template size encompasses all pixels that could be reached given the imaging field of view size (*X* × *Y*) and the StripRegistration parameter `maxShift` (*m*). This padded image is the initial template, *T*(1) ∈ ℝ^(*X*+2*m*)×(*Y*+2*m*)^.

Then, the motion displacement, (*s*_*x*_ (*t*), *s*_*y*_ (*t*)) at time point *t*, of the corresponding frame of the movie, *F*(*t*) ∈ ℝ^*X*×*Y*^, is estimated by finding the shift that had the highest pixel-value correlation to a dynamic template 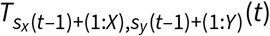 at that point in time, cropped to the best aligned imaging field of view from the previous time point. NaN values in the template are filled in with 0 for this calculation. At each frame, the algorithm searches over displacements that are within the StripRegistration parameter `clipShift` (*c*) pixels from the estimated motion at the previous timestep:

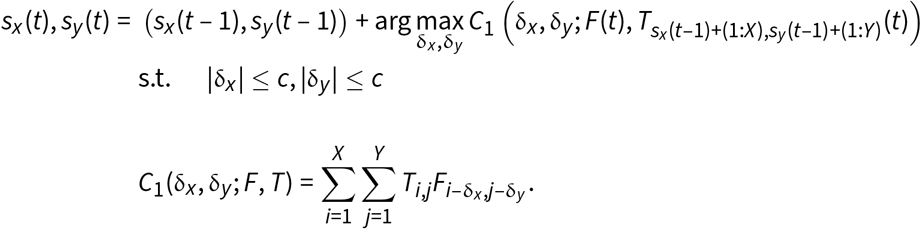

This optimization problem is solved using a matrix-multiply discrete Fourier transform approach, which also allows for estimation of subpixel shifts (Guizar-Sicairos et al. 2008). We use this to refine our motion estimate to quarter-pixel precision.

If the magnitude of the estimated motion was large 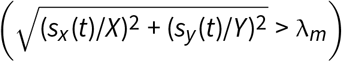, we re-solve for *s*_*x*_ (*t*), *s*_*y*_ (*t*), using a normalized cross-correlation measure that operates only on the set of pixels (Ω) that observed in the data *F*(*t*) and that are valid (not NaN) for all possible shifts of the template. We relax the constraints on δ_*x*_ and δ_*y*_ to span a –50 to 50 pixel range of motion displacements around the previous motion estimate, resulting in the optimization

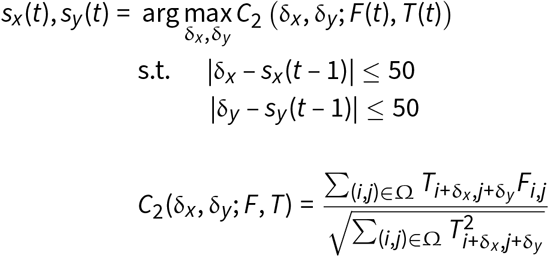

To estimate a subpixel motion shift with this metric, the values of *C*_2_ around the maximum were assumed to be quadratic along each the *x* and *y* axes, and the peak of this quadratic was taken as the subpixel motion shift. A threshold around λ_*m*_ = 0.6 worked well, and we found that this re-optimization step provided more accurate estimates when the motion was large relative to the imaging FOV and prevented noisy frames from causing the motion estimates to drift away from the global optima. For both estimation methods, motion estimates that were outside the bounds dictated by *m* along either axis were rejected.

Once the motion displacement is estimated, we update the dynamic template by averaging the initial template, which is centered on the dynamic template canvas, with pixel values that had already been observed at least 100 times from already motion corrected frames. Mathematically we write

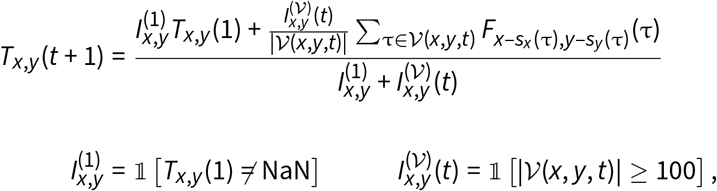

where *V*(*x, y, t*) is the set of time points less than *t* during which pixel (*x, y*) was observed after motion correction and 1[·] is the indicator function that evaluates to 1 if its argument is true and 0 if not. For calculations, NaN values in *T*_*x,y*_ (1) are ignored.

The final set motion estimates over time are generated by repeating the above steps sequentially through all time points. If the data were downsampled, the motion estimates are resampled to the original temporal resolution using piecewise cubic hermite interpolating polynomial (PCHIP) spline interpolation. A motion corrected movie 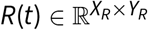 is also produced by shifting each raw frame or downsampled frame *F*(*t*) within the (*X* + 2*m*) × (*Y* + 2*m*) expanded field of view and cropping to the central *X*_*R*_ × *Y*_*R*_ pixels, where *X*_*R*_ = *X* + 2 max_*t*_ |*s*_*x*_ (*t*)| and *Y*_*R*_ = *Y* + 2 max_*t*_ |*s*_*y*_ (*t*)|. Unobserved pixels in this expanded field of view at each frame are padded with NaNs.

After motion correction, for each downsampled time point, we calculate a rectified negative error (RecNegErr) metric per registered frame, *R*(*t*), relative to the median of nearby registered frames 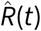. The RecNegErr calculation is performed in 10 s chunks of time and 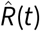 is calculated over that chunk as

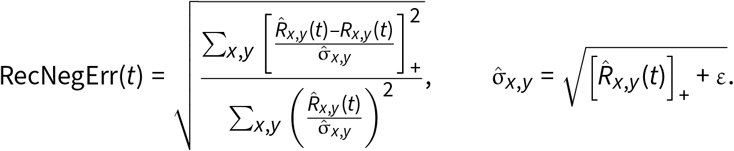

This metric was used to censor frames that had excessive out-of-plane motion in downstream analyses. In simulations, RecNegErr on frames was correlated with its simulated out-of-plane motion displacement (Figure S2).

#### Other motion correction methods

To compare StripRegistration against existing methods, we ran our data through NoRMCorre (Pnevmatikakis & Giovannucci 2017) using the CaImAn package (Giovannucci et al. 2019) and the motion correction step of Suite2p (Stringer et al. 2026). We used as large of a maximum displacement parameter as each algorithm would allow in order to give each motion correction method a fair chance to correct large motion deviations relative to the imaging field of view size.

#### Source extraction with SILo

Source Identification by Activity Localization (SILo) is inspired by super-resolution localization microscopy methods (Lelek et al. 2021) and comprises event detection, source localization, and fluorescence trace extraction. We perform event detection and source localization in the downsampled temporal space, which has *N*_*ds*_ frames, and we perform fluorescence trace extraction at the full temporal resolution, which has *N*_*t*_ = 2^*d*^*N*_*ds*_ frames.

For SILo, we first want to censor frames with residual motion (e.g. out-of-plane motion) using a high pass filtered and normalized version of RecNegErr, ε(*t*). The set of misaligned frames (*D*) is

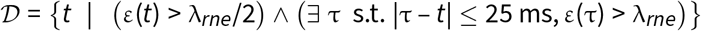

where λ_*rne*_ is the SILo parameter `motionThresh`.

Then, the set of valid, well-aligned frames to keep for downstream analyses, T is

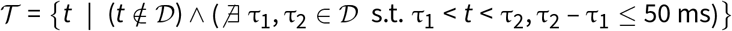

All pixels within frames not in *T* were set to NaN (i.e. *R*_*x,y*_ (*t*) = NaN, ∀ (*x, y*), *t* ∉ *T*). Along the spatial dimensions, we define a set of valid pixels for further analysis as

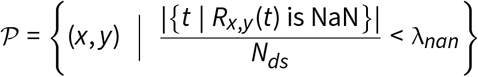

where λ_*nan*_ is the SILo parameter `nanThresh`.

Event detection: To stabilize the statistics of each pixel for event detection, we first estimate the baseline 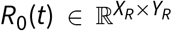 and standard deviation 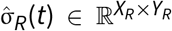 for every valid pixel within every frame. 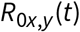 at each pixel is estimated by a moving median filter over time on a slightly temporally smoothed version of *R*_*x,y*_ (*t*) with a window length of the SILo parameter `baselineWindow_Glu_s`. The smoothing is performed with a moving mean over time of window length `denoiseWindow_s`, which makes the median filter less sensitive to discrete photon counts in dim pixels. The standard deviation was estimated using a linear model as

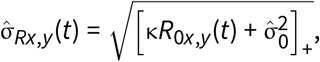

where κ is the multiplicative relationship between mean and variance, and 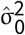 is the estimated variance of the noise floor. These parameters are estimated using the variances and means of valid pixels over the first 500 valid frames. 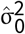 is estimated as the 10^th^ percentile of those variances, scaled by a variance inflation factor specified by the SILo parameter `VIF`, and κ is estimated using the ratio of the variances and means after accounting for the noise floor. We then calculate a baseline-subtracted and standardized version of the data:

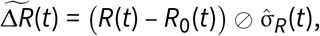

where ⊘ represents element-wise division.

To filter for glutamate indicator fluorescence events in the data, 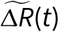 is convolved in space with a difference of Gaussians kernel (σ_1_ = `sigma_px` pixels, σ_2_ = 5σ_1_) and in time with a exponential (τ =`tau_s`). This step serves as a “bump detector” where the negative dip surrounding the central Gaussian bump prevents flat areas of fluorescence from triggering false positives. Mathematically we write

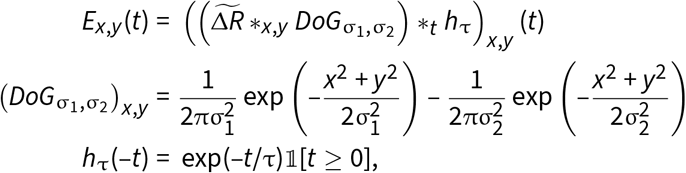

where *_*x,y*_ and *_*t*_ indicate convolutions over the spatial and temporal axes, respectively.

Local maxima in space and time of *E*(*t*) are considered “candidate events” and are used to generate an activity image, 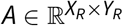 computed as

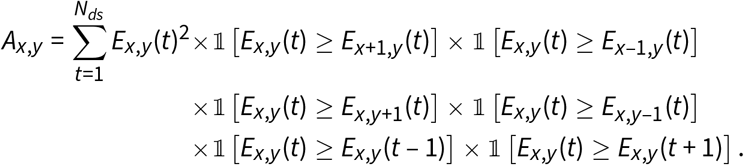

The final SILo activity image, 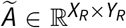, is a spatially high pass filtered version of *A* to reduce low frequency features in the activity image, which do not represent localized glutamate indicator fluorescence transients.

Source localization: Source localization is performed using the SILo activity image by fitting Gaussians to the SILo activity image. This step is motivated by the fact that isotropic deviations of centroid locations of single sources will cluster around the actual position of the source. Thus we model *Ã* as the sum of Gaussians, each corresponding to a localized source:

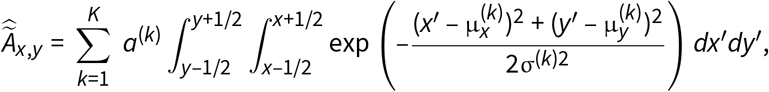

where *K* is the total number of sources, and for the *k*^th^ source, *a*^(*k*)^ is the amplitude of its peak, 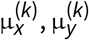 is the coordinate of the center of the peak, and σ^(*k*)2^ is the variance of the activity profile. *K* is estimated iteratively as follows. The baseline intensity, *b*, and noise, σ_*b*_, of the *Ã* are estimated as the median and robust standard deviation estimate over pixels, respectively. Then, the all local maxima in *Ã* with a neighbor with an intensity larger than *b* + λ_*p*_σ_*b*_ are seeded as sources, where λ_*p*_ is the SILo parameter `peakth`. These local maxima sit on the integer pixel grid and are denoted 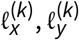 for the *K*_0_ initial sources. The parameters of these peaks is then optimized by

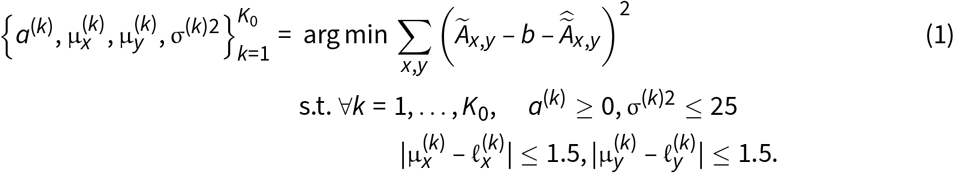

With the initial set of *K*_0_ peaks we then iteratively add new peaks to ensure that we capture the full set of sources in the data. The next candidate peak is given by

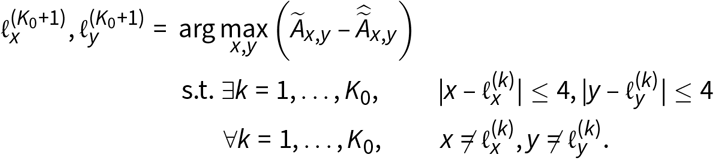

If 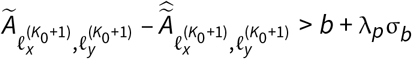, this source is added to the set and the parameters of the new set of peaks are optimized using Equation 1 and the iterative process continues. If the peak is not large enough, the algorithm stops searching for more sources to add.

Finally, we filter out peaks that are likely to be spurious: any peak that has an low amplitude relative to its spread, specifically if 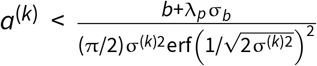, is removed. After filtering, we update the number of sources identified as *K*.

Fluorescence trace extraction: The final set of source locations is then used to seed the fluorescence trace extraction step. Because we wish to estimate the fluorescence traces the imaging rate, this step takes the motion corrected data 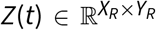 at the full temporal resolution as input. The set of invalid frames is upsampled to the full temporal resolution as well by setting a frame to be invalid if either of its neighboring downsampled time points is invalid.

To account for the noise statistics of the data, we convert the units of the data into photon counts, which are Poisson distributed, by taking 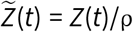, where ρ is the SILo parameter `photonScale`. If ρ is not provided by the user, we estimate the photon scale using the ratio of variance and means of valid pixels after accounting for the noise floor 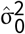.

With 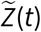, we then solve a constrained nonnegative matrix factorization problem. To assist with notation, we rearrange the data as 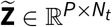, where *P* = |*P*| is the total number of valid pixels in the data and each column of 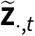 contains the values of each valid pixel at frame *t*. The spatial profiles of sources, **S** ∈ ℝ^*P*×*K*^, the temporal traces of sources, 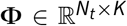, and a time varying background, 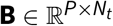, are fit by the following constrained optimization

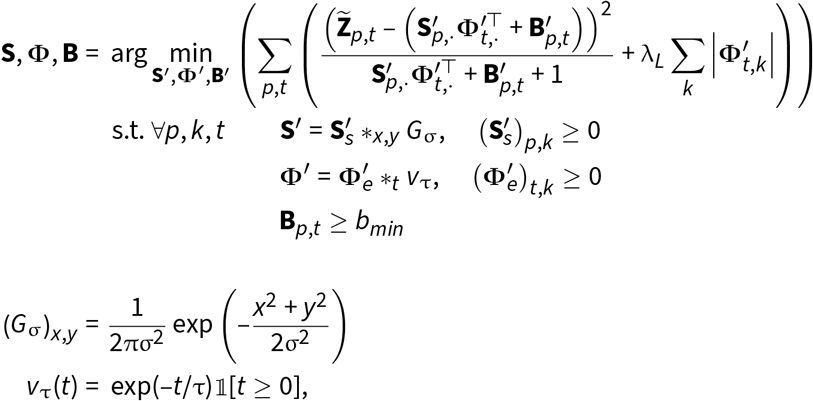

where *b*_*min*_ is the SILo parameter `minBaseline` and λ_*L*_ (*ℓ*_1_-regularization coefficient) is the SILo parameter `lambda`. The optimization is solved by an iterative coordinate descent over `nmfIter` iterations, which we set as a SILo parameter. After fitting, a debiasing step is performed by refitting **Φ** using the same objective, but restricting the support of 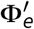 to that of the original fit and replacing the *ℓ*_1_-regularization coefficient with ϕ · λ_*L*_, where ϕ is the SILo parameter `phi`. The debiased **Φ** and optimized **S** are the main outputs of SILo.

In addition to returning **Φ**, which is a denoised version of the fluorescence traces, we also return an estimate of the fluorescence traces from a least squares fit of the spatial profiles to the data:

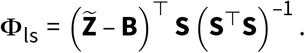

#### Other source extraction methods

We compare SILo against existing methods, running our data through CNMF (Pnevmatikakis et al. 2016) using the CaImAn package (Giovannucci et al. 2019) and Suite2p (Stringer et al. 2026). Because we found that StripRegistration gave the fewest errors in motion correction for our data, we provided all source extraction algorithms with the StripRegistration-corrected data. As the motion sometimes causes pixels in the movie to become unobserved, we cropped the motion corrected movie to only pixels that were observed at least 35% of the time for CaImAn and to pixels that were observed at least 94% of the time for Suite2p. Missing pixel values were filled in by linearly interpolating in time. To fairly compare between source extraction methods, we optimized parameters for SILo, Suite2p, and CaImAn on the “default” simulations (motion standard deviation of 1 µm, brightness of 0.6 photons, and 30 sources) using Bayesian optimization (Frazier 2018). These parameters were then fixed when evaluating source extraction across all simulation conditions.

### Evaluation metrics

#### Motion correction

In simulated data, estimated motion vectors were compared to ground truth motion vectors using the root mean squared error (RMSE)

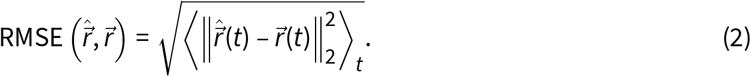

For *in vivo* data, a goodness-of-alignment score was calculated using the motion corrected movie. We used the mean frame correlation to the mean image (mCM) and the correlation between the mean image and maximum image (CMM) (Hattori & Komiyama 2022). To reduce noise in the maximum projection image, the movie was first downsampled by averaging every four frames before calculating the maximum of each pixel across time.

#### Source extraction

In simulated data, the source detection performance was quantified by the *F*_1_ score relative to the ground truth sources. *F*_1_ is calculated from the number of true positives (TP), false positives (FP), and false negatives (FN):

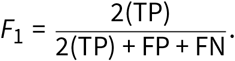

To assign extracted sources as TPs, a one-to-one matching was performed by solving a linear sum assignment problem, where the cost was related to the cosine similarity between the extracted source spatial profile and the ground truth spatial profile. Matches were also constrained to require a cosine similarity of at least 0.5.

For fluorescence trace extraction, we compared the ability of each method to demix overlapping sources by calculating a “demixing purity” metric. For an extracted TP source *k* with fluorescence trace 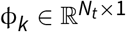 and *M* spatially overlapping ground truth sources with fluorescence traces 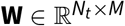, we fit

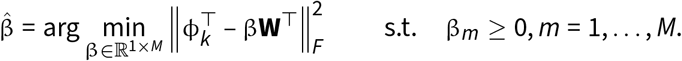

By convention, we let the ground truth trace that is matched to the extracted TP source be the first column of **W**. The demixing purity for source *k*, η_*k*_ is then given by

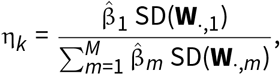

where SD(·) is the standard deviation of the elements of the provided vector. As η_*k*_ represents how concentrated the representation of the extracted trace is on the ground truth trace, a value of 1 means that the extracted trace is purely representing the GT trace. A low value would mean that there is significant activity not representative of the GT trace but instead representative of other traces, indicating bleedthorugh.

For an extracted fluorescence trace, ϕ(*t*), we calculated peak SNR as the ratio of the 99^th^ percentile of ϕ(*t*) to an estimate of the baseline noise in the trace, based on the standard deviation of the least squares estimate of the trace, ϕ_ls_(*t*), when ϕ(*t*) < ε, for small ε:

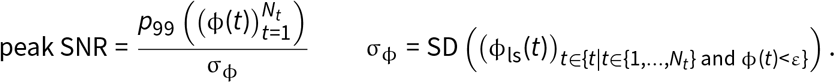

### Retrospective ex vivo synapse localization

#### Histology sample preparation

The same day after each mouse underwent *i*n vivo glutamate imaging, each mouse was anesthetized with an overdose of isoflurane and transcardially perfused first with 10 mL 1x PBS (flow rate 7 mL/min) and then 50 mL 4% paraformaldehyde and 10% acrylamide in 1x PBS (flow rate 7 mL/min). Each brain was sliced into 100 µm thick slices in a plane parallel to the cranial window using a similar procedure as Ouellette et al. 2022. The slice with the dendrites that were imaged *in vivo* were identified by imaging the slices on a spinning disk microscope, and the slice was trimmed to the area containing the dendrites. This trimmed slice was then prepared into a gel using the LICONN protocol (Tavakoli et al. 2025). Gels were labeled with the antibodies listed in Table 1.

**Table 1:** Antibodies used for immunofluorescence.

| Antibody Target | Fluorophore | Species | Vendor | Catalog # |
| --- | --- | --- | --- | --- |
| Bassoon | – | Rabbit | Synaptic Systems | 141 003 |
| PSD-95 | – | Mouse (IgG2a) | Antibodies, Inc. | 75-028 |
| Gephyrin | – | Mouse (IgG1) | Synaptic Systems | 147 111 |
| Green fluorescent protein (GFP) | Dylight™ 405 | Goat | Rockland | 600-146-215 |
| Mouse IgG1 | ATTO 488 | Goat | Rockland | 610-152-040 |
| Rabbit IgG (H&L) | ATTO 647N | Goat | Rockland | 611-156-122 |
| Mouse IgG2a | ATTO 550 | Goat | Rockland | 610-154-041 |

For imaging, gels were expanded in purified water (Millipore Milli-Q) overnight, mounted on a poly-lysine coated holder, and imaged using a Zeiss Lightsheet 7 microscope using a ×20, 1.0NA water immersion objective.

#### Analysis of structural images

To preliminarily align each histology image to the corresponding *in vivo* two-photon image, matching key points were manually selected on both the maximum projection over the depth axis of the iGluSnFR channel of the histology image and the average image over time of the two-photon recording. Using these key points, the histology image was aligned into the space of the two-photon image using a 2D projective geometric transform, that was applied to each plane of the histology volume. With this preliminary alignment, the centroids of the sources detected by SILo from the two-photon recording were mapped into the histology volume. Additional control centroids (equal in number to the number of detected sources) were dropped randomly in other areas of bright fluorescence to maximize the total coverage of control centroids and source centroids along the imaged segment of dendrite. All synapses around each centroid were then manually annotated, with the annotator blind to whether the centroid was of a SILo-detected source or a control location. Synapses were defined as a region of iGluSnFR-labeled membrane coincident with a patch of labeled Bassoon and labeled PSD-95. Neighboring patches were labeled as distinct synapses if both the Bassoon patches and PSD-95 patches were discontinuous between synapses.

With the set of annotated synapses on the histology data and the set of SILo-detected sources, sources and synapses were matched by solving a linear assignment problem, where the cost was the 2D distance between the centroids of elements of a matched pair and the cost of being unmatched was around 0.8 µm. “Near matches” were defined as sources that were unmatched by this problem but were still within 0.75 µm of an annotated synapse. To display the final SILo-detected source profiles on the histology images, a final non-rigid spline alignment was performed with the centroids of matched synapses and SILo-detected sources to place the *in vivo* two-photon image into the space of the histology image. We quantified matching using using the positive predictive value (PPV):

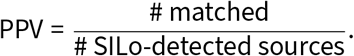

## Data and code availability

MATLAB code for GIAnT is publicly available through GitHub (https://github.com/AllenNeuralDynamics/GIAnT-MATLAB), and a Python implementation is forthcoming. Experimental data is available available on the `aind-open-data` S3 bucket (MISSING).

## Acknowledgments

We thank the Lab Animal Services, Neurosurgery & Behavior, Scientific Instrumention & Process Engineering teams at the Allen Institute for technical support, K. Cao and J. Chandrashekar for helpful discussions on the expansion microscopy and immunolabeling protocol, B. Cruz for assistance developing Bonsai workflows, D. Birman and S. de Vries for help organizing our data for release, and T. Zeric and K. Girven for project management support.

This research was supported by the Allen Institute, founded by Jody Allen — chair and co-founder of Allen Family Philanthropies, and the late Paul G. Allen — investor, philanthropist, and co-founder of Microsoft. We gratefully acknowledge their vision and generosity, which make this work possible. This research was also supported by the Chan Zuckerberg Initiative (CP2-1-0000000704) and National Institutes of Health (NINDS: DP2NS136990, NIBIB: R01EB037653, NIMH: F30MH138009, and NIGMS: T32GM136577). The content is solely the responsibility of the authors and does not necessarily represent the official views of the National Institutes of Health. MEX was supported by the Paul and Daisy Soros Fellowships for New Americans.

## Author contribution matrix

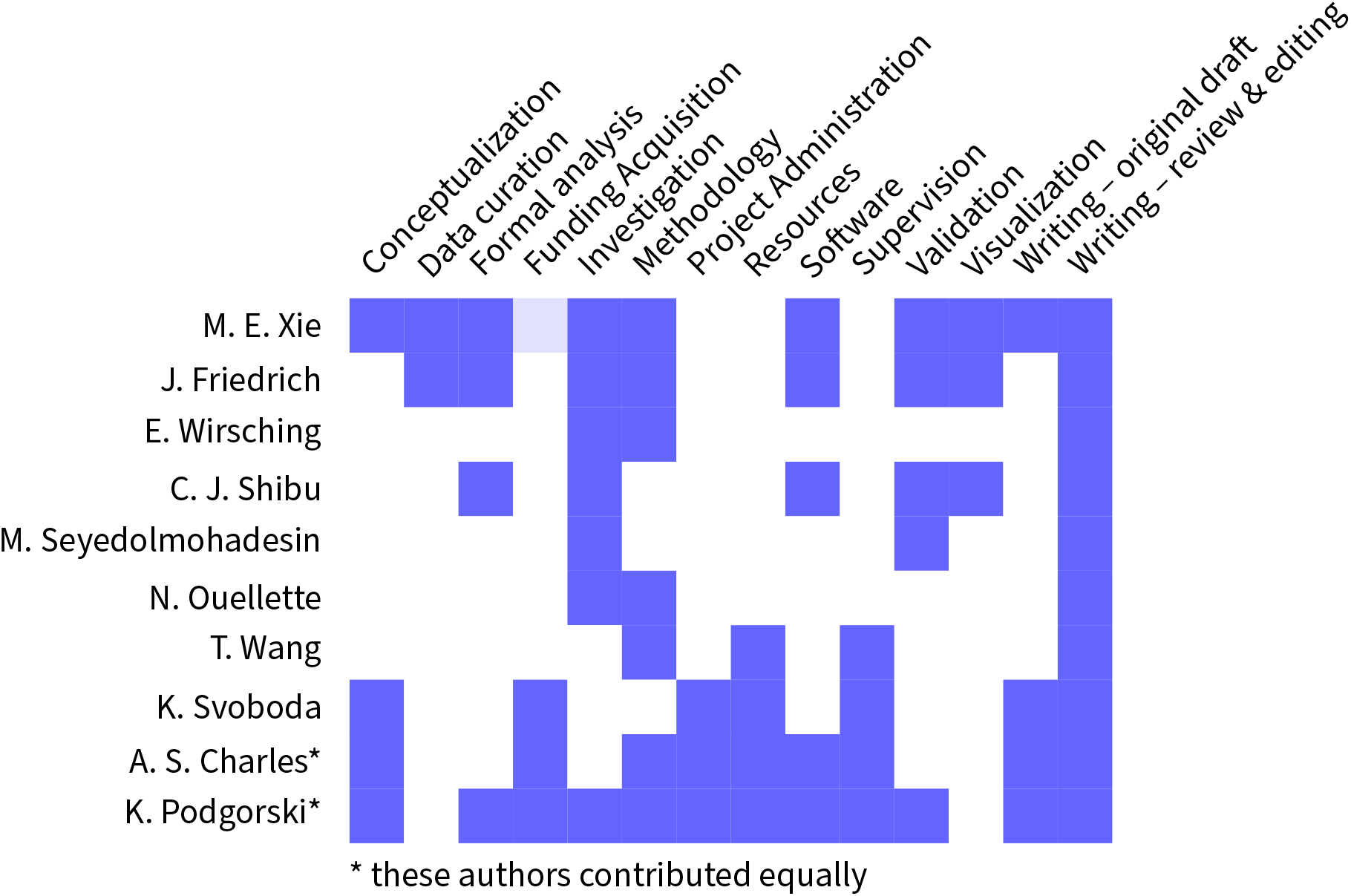

---

**Figure S1:**
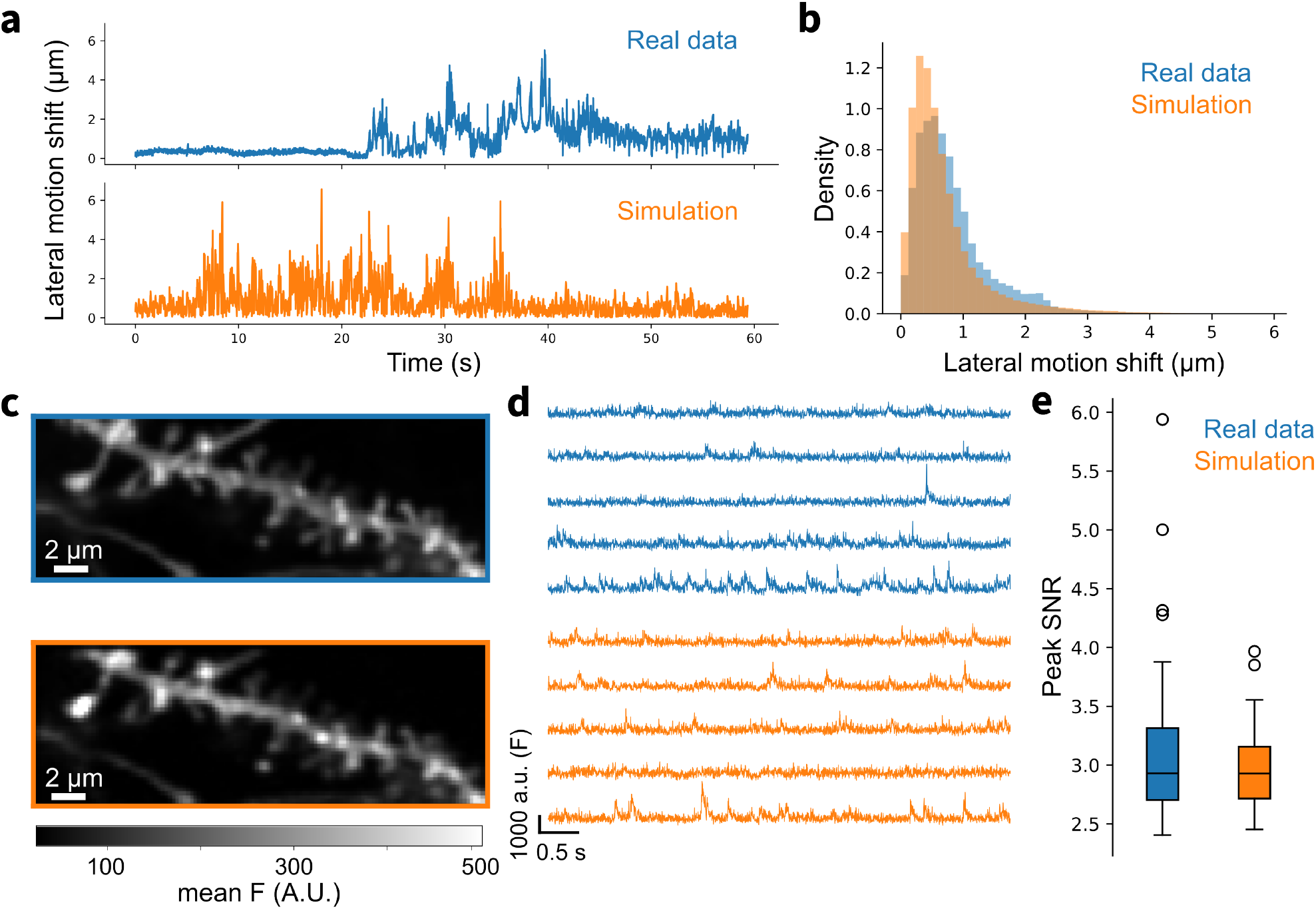
Statistics of default simulations (motion σ=1 µm, brightness=0.6 photons, 30 sources) compared to real *in vivo* two-photon imaging data. **a**. Lateral motion shift estimated from real data using StripRegistration (blue) and ground truth lateral motion shift used to generate simulated data (orange). **b**. Distribution of estimated lateral motion shift from real data (blue) and ground truth lateral motion shift for simulations (orange) across 12 datasets of each. **c**. Mean fluorescence over time of motion corrected real data (blue) and simulated data (orange). **d**. Fluorescence traces from real data (blue) and simulated data (orange), generated by averaging the nine pixels around the five largest peaks in the variance image. **e**. Distribution of peak signal-to-noise ratio (SNR) of the five pixel-averaged fluorescence traces from 12 real data sets (blue) and simulated data sets (orange).

**Figure S2:**
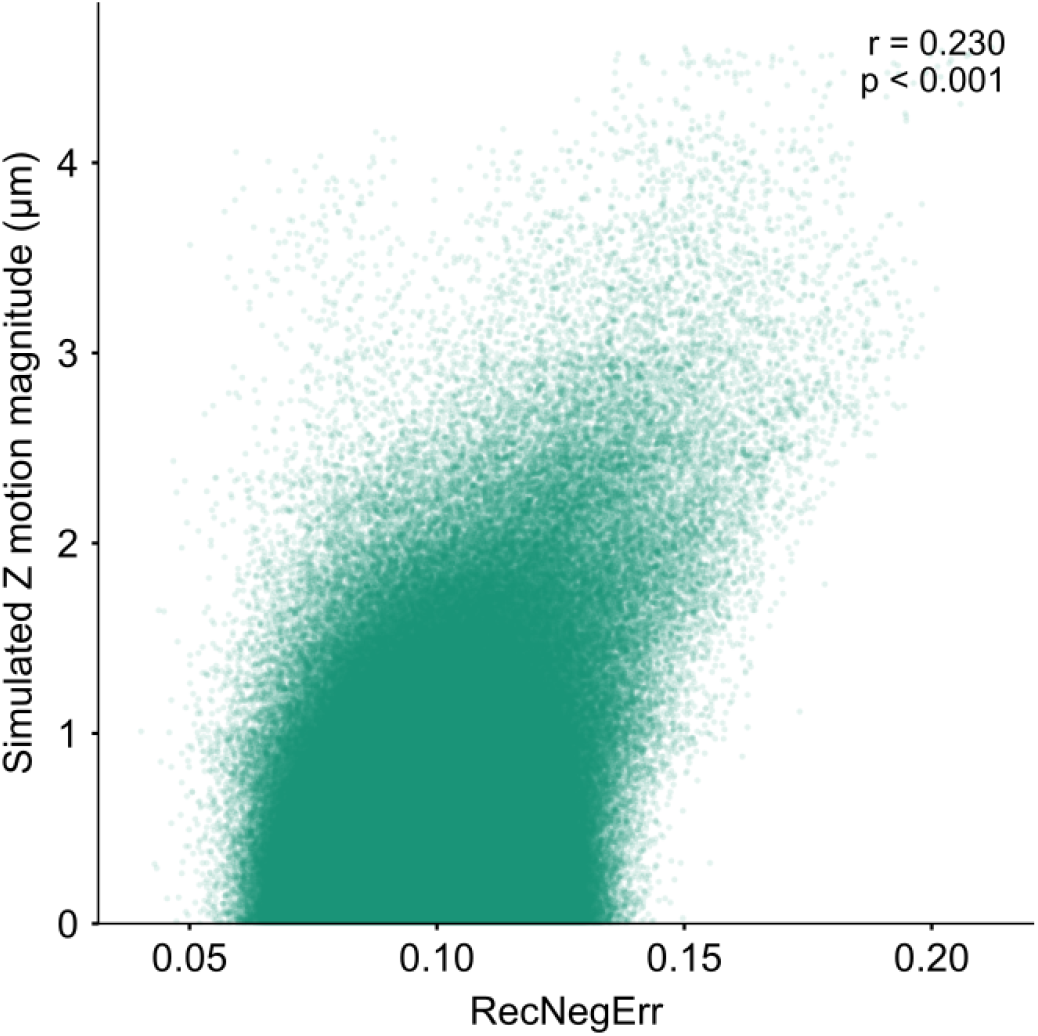
Rectified negative error (RecNegErr) vs. out-of-plane motion in simulations. Scatterplot of RecNegErr versus ground truth out-of-plane (Z) motion magnitude. Each point represents one frame within a default simulation, and the plot is pooled across all 12 default simulations.

**Movie S1:**
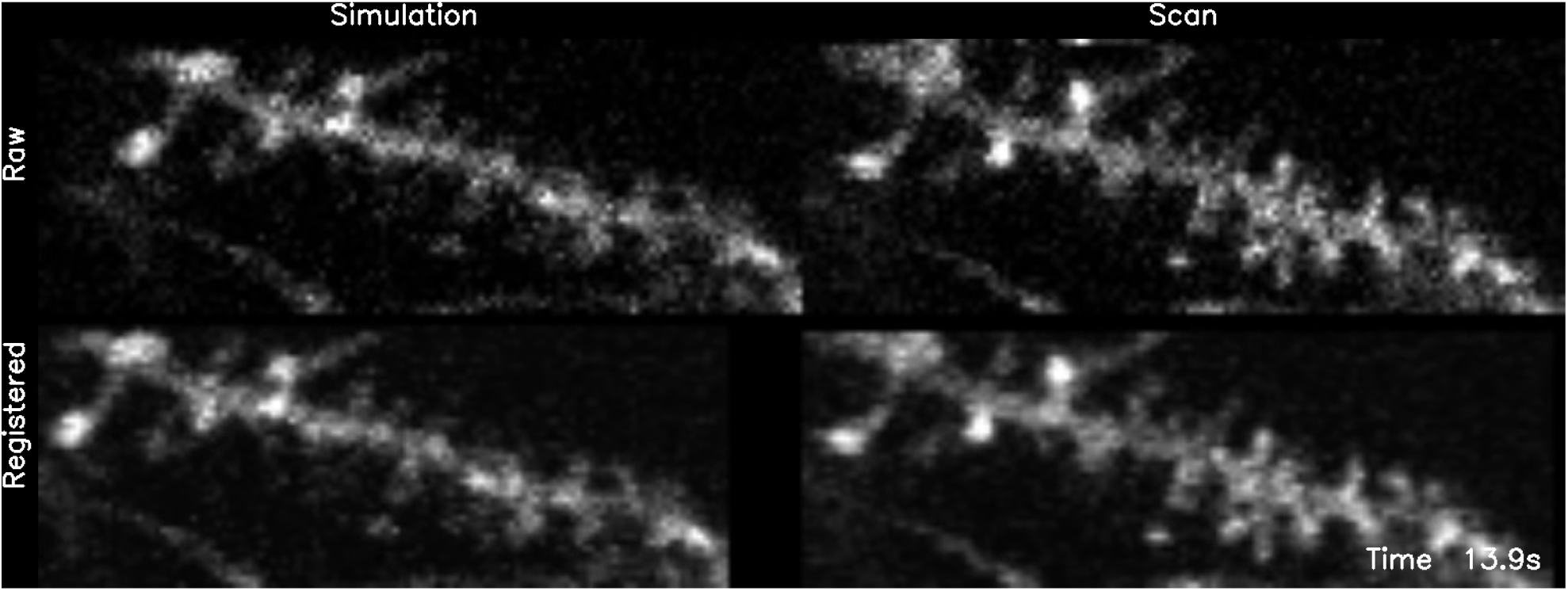
StripRegistration on simulations and *in vivo* scans. Raw simulations and *in vivo* data (top left and right, respectively) were motion corrected with StripRegistration (bottom left and right). For the movie visualization, movies are temporally downsampled 20×.

**Movie S2:**
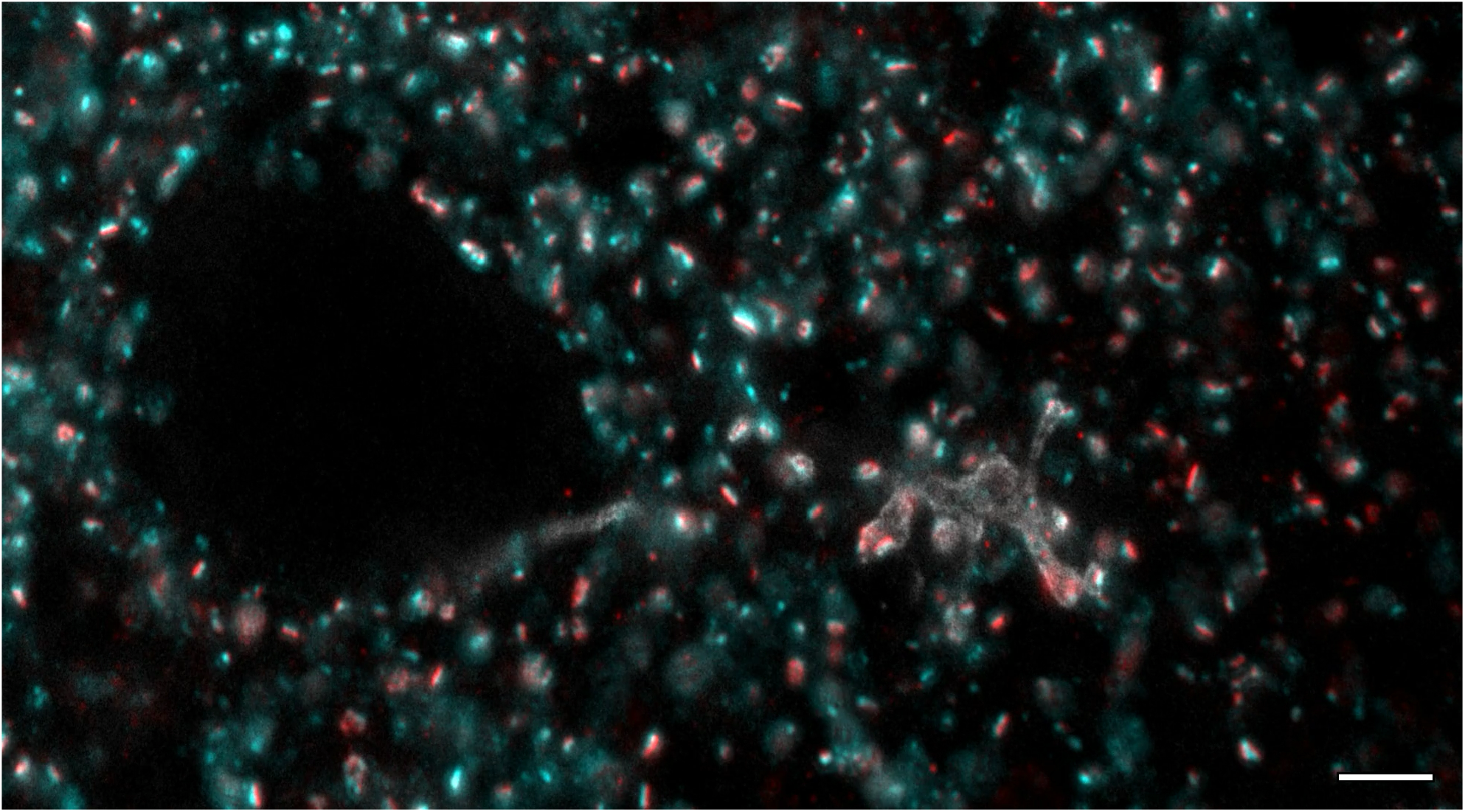
Histology volume of an example dendrite after immunolabeling and expansion. Expanded volume of dendrite immunolabeled for iGluSnFR (white), Bassoon (cyan), and PSD-95 (red). Each frame of the movie is a different plane of the volume. Scale bar, ∼2 µm before expansion.

